# Bacterial produce membrane-binding small molecules to regulate horizontal gene transfer in vesicles

**DOI:** 10.1101/2022.04.08.487716

**Authors:** Frances Tran, Manasi S. Gangan, James Q. Boedicker

## Abstract

The exchange of bacterial extracellular vesicles facilitates molecular exchange between cells, including the horizontal transfer of genetic material. Given the implications of such transfer events on cell physiology and adaptation, some bacterial cells have likely evolved mechanisms to regulate vesicle exchange. Past work has identified mechanisms that regulate the formation of extracellular vesicles, including the production of small molecules that modulate membrane structure, however whether these mechanisms also regulate vesicle uptake and have an overall impact on the rate of vesicle exchange is unknown. Here we show that membrane-binding molecules produced by microbes regulate both the formation and uptake of extracellular vesicles and have the overall impact of increasing the vesicle exchange rate within a bacterial coculture. In effect, production of compounds that influence vesicle exchange rates enable cells to steal genes from neighboring cells. The ability of several membrane-binding compounds to regulate vesicle exchange was demonstrated. Three of these compounds, nisin, colistin, and polymyxin B, are antimicrobial peptides added at sub-inhibitory concentrations. These results suggest that a key function of exogenous compounds that bind to membranes may be the regulation of vesicle exchange between cells.

**Importance:** The exchange of bacterial extracellular vesicles is one route of gene transfer between bacteria, although it was unclear if bacteria developed strategies to regulate the rate of gene transfer within vesicles. In eukaryotes, there are many examples of specialized molecules that have evolved to facilitate the production, loading, and uptake of vesicles. Recent work with bacteria has shown that some small molecules influence membrane curvature and induce vesicle formation. Here we show that similar compounds facilitate vesicle uptake, thereby regulating the overall rate of vesicle exchange within bacterial populations. The addition of membrane-binding compounds, several of them antibiotics at sub-inhibitory concentrations, to a bacterial co-culture increased the rate of horizontal gene transfer via vesicle exchange.

## Introduction

Many biomolecules are exchanged via bacterial extracellular vesicles. Bacterial vesicles are known to contain cytoplasmic and membrane proteins, genetic material, and small molecules including bacterial signaling molecules. The uptake of vesicles enables molecular transfer between different species of bacteria and from bacteria to eukaryotic host cells (1-6). Vesicle exchange contributes to horizontal gene transfer within bacterial populations (7-10). Although many mechanisms have been shown to regulate bacterial vesicle formation (11-19), less is known about mechanisms cells use to control the exchange of vesicles, which involves both the production of vesicles by a donor cell and the uptake of vesicles by a recipient cell. Given the importance of vesicle exchange to many cellular processes and the ubiquity of vesicle production by many bacterial species (20, 21), it seems likely that bacteria would have evolved strategies to elicit and control vesicle exchange.

More is known about regulation of vesicle exchange within eukaryotic systems. Eukaryotic vesicles are essential to signal transmission within neuronal synapses and also involved in immune regulation and angiogenesis (22-26). Vesicle formation and uptake both require restructuring the membrane and the formation of energetically costly intermediate states of the membrane (27-29). Eukaryotic cells overcome these energy barriers through the use of molecular motors and membrane-restructuring molecules to induce membrane curvature (30-32). Similar strategies have been shown in bacteria, with the best example being regulation of vesicle production via pseudomonas quinolone signal (PQS) (11). PQS inserts into the bacterial membrane, inducing curvature and leading to increased vesicle production (11, 14, 33). PQS production can also induce vesicle formation in neighboring species (5). Other membrane-binding compounds have been shown to influence vesicle production, including polymyxin B, colistin, and phenol-soluble modulins (34-36). These reports show that as in eukaryotic cells, vesicle production by bacteria can be regulated by molecules that bind to and restructure the cell membrane.

It is not known if molecules that restructure the cell membrane also influence vesicle uptake by bacteria, and if the presence of such molecules impacts the overall rate of vesicle exchange within a population of bacteria. Here we test the influence of several membrane-restructuring compounds on the rate of vesicle production and vesicle uptake to determine the extent that vesicle exchange can be regulated via exogenous compounds. Vesicle uptake was quantified through the vesicle-mediated transfer of plasmid DNA and the resulting gain of antibiotic resistance in the recipient population (10). These results demonstrate that exogenous bacterial compounds that are known to bind to and restructure the cell membrane regulate vesicle exchange within bacterial populations.

## Results

### Membrane structuring protein alpha-synuclein regulates the production and uptake of extracellular vesicles in bacteria

In eukaryotic systems, the production and uptake of vesicles is regulated by many mechanisms. One mechanism for EV biogenesis in eukaryotic systems includes recruitment of ESCRT (endosomal sorting complexes required for transport) complexes and their interaction with the membrane and many other factors (37, 38). As for EV uptake in eukaryotic systems, EV binding and uptake can be regulated by transmitted signals from the cell surface to elicit uptake (39). As vesicle exchange in bacterial cells could also involve restructuring and reshaping the cell membrane, we sought to determine if biomolecules known to interact with cell membrane would regulate exchange of bacterial vesicles. Initial experiments examined the influence of the well-characterized human protein, alpha-synuclein (AS), on vesicle formation and uptake. AS binds to membranes and is found in high abundance in presynaptic terminal associated with synaptic vesicles (40-42). Alpha-synuclein binds to curved, anionic lipids (43). In addition, previous studies have suggested membranolytic effect of AS on bacterial cell (44). We speculated that the ability of AS to bind to and restructure cellular membranes, would translate to modulation of vesicle production and uptake in bacteria at sublethal concentrations.

To test the ability of AS to influence vesicle production, concentrations of purified AS between 0.01 µM and 0.1 µM were added to cultures of *Escherichia coli* MG1655 containing the plasmid pLC-RK2 (10), see Table S1. Vesicles were harvested from culture after 16-20 h of growth via size-exclusion filtration and ultra-centrifugation, see Fig 1A. Production of vesicles was measured by quantifying the concentration of outer-membrane proteins, OmpC/F, in solutions of harvested vesicles via SDS-Polyacrylamide gel electrophoresis, see Fig S1. As shown in Fig 1B, cultures of the *E. coli* donor strain grown in AS resulted in 2 to 3 times more vesicle production. AS at 0.1 µM did not strongly influence cell growth, see Fig S2.

**Figure 1:**
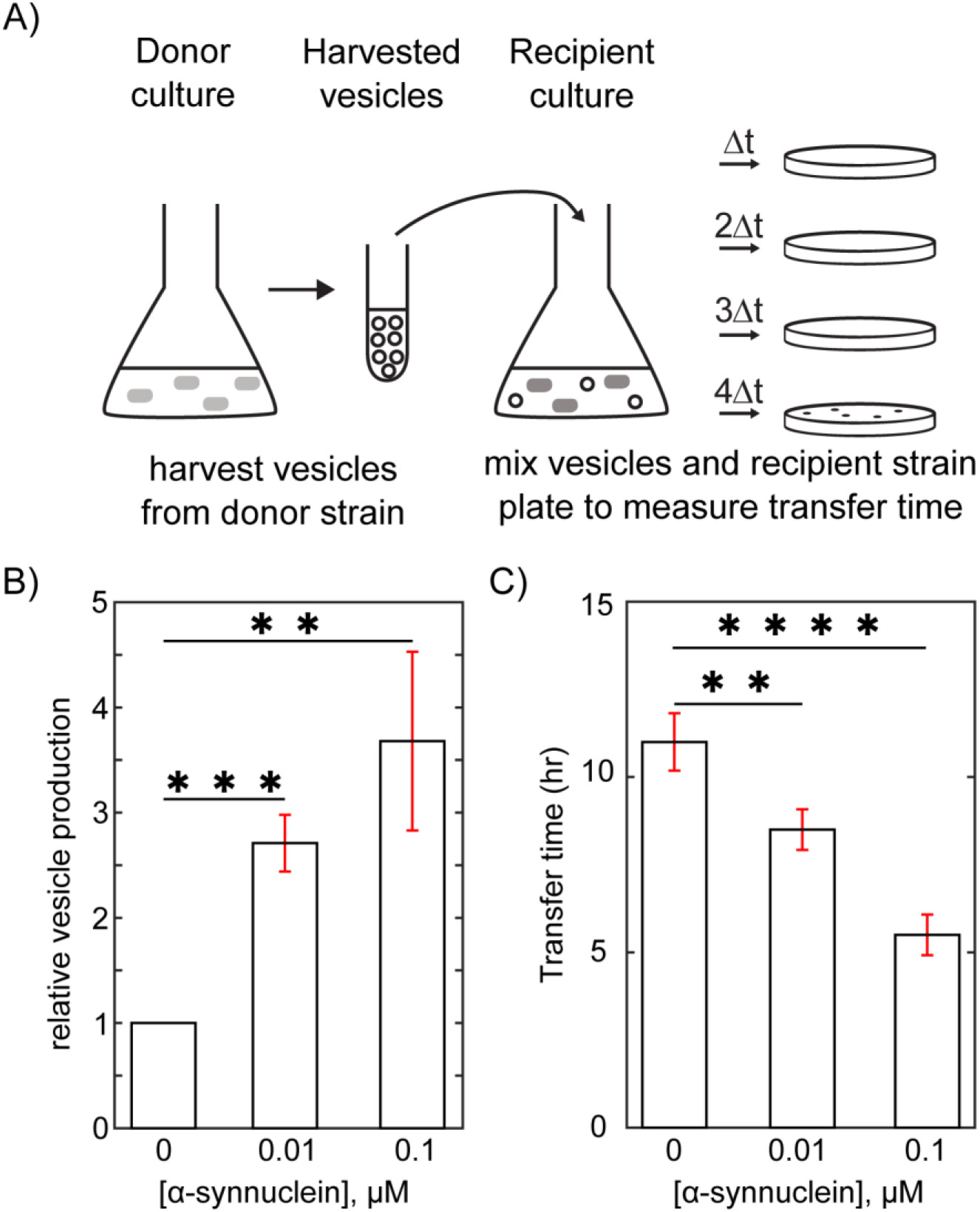
Alpha-synuclein increases the rates of extracellular vesicle (EV) production and uptake. A) EVs were harvested from a donor culture via filtration and centrifugation. The donor strain contained a plasmid conferring antibiotic resistance. Harvested Evs were added to a recipient culture, and EV uptake was monitored by detecting the gain of resistance in recipient cells. B) Addition of the membrane binding eukaryotic peptide alpha-synuclein increased the rate of vesicle production by the *E. coli* donor strain in a dose-dependent manner. C) Addition of alpha-synuclein to the recipient *E. coli* culture decreased the time to transfer of EVs in a dose-dependent manner. n=3. Error bars show standard deviation. Significance in the difference observed in vesicle production and transfer time between treated and untreated samples was confirmed with unpaired t test (** P ≤ 0.01, *** P ≤ 0.001, **** P ≤ 0.0001).

Next, we tested if these same concentrations of AS would likewise influence the uptake of vesicles by a recipient strain. The assay for vesicle uptake is depicted in Fig 1A. Vesicles were harvested from a donor bacterial strain containing a plasmid, and the harvested vesicles, some containing the plasmid pLC-RK2, were added to a recipient bacterial strain. Aliquots of the culture of receiver strain with added harvested vesicles were removed at a set time interval and spread onto antibiotic selection plates. The plasmid contained a resistance marker, and the recipient strain did not grow on antibiotic selective plates in the absence of the plasmid. The time needed to detect a recipient cell with antibiotic resistance was defined as the time to transfer and is proportional to the rate of successful gene transfer via vesicles. In previous studies, we have shown that gene transfer in vesicles has a characteristic transfer time that depends on the concentrations and characteristics of the transferred plasmid, the donor strain, and the recipient strain (10). Gain of resistance in this assay is the result of the uptake of plasmids located inside of harvested vesicles, as verified by detection of the transferred plasmid in resistant recipient strains via colony PCR. Vesicles from an *E. coli* MG1655 donor strain containing plasmid pLC-RK2 were added to the recipient strain, *E. coli* MG1655, at early exponential growth phase. In transfer experiments a standard number of vesicles was used. Vesicles added to recipient cultures contained a total of 1 µg of the outer membrane proteins OmpC/F, quantified via protein gels, see Fig S1. As shown in Fig 1C, in the absence of AS gene transfer occurred after 11 h, whereas the time to transfer was shortened to 8.5 and 5.5 h after adding 0.01 µM and 0.1 µM AS, respectively.

Increased vesicle production and uptake rate in the presence of AS suggested that exogenous molecules known to bind to, and restructure cellular membranes have the potential to modulate vesicle exchange between bacterial cells. Next, we tested if this phenomenon was general to other exogenous biomolecules known to interact with outer membranes, specifically compounds naturally released by bacteria.

### Membrane binding exogenous molecules produced by bacteria increased vesicle production

Many molecules released by bacteria are known to bind to and restructure cellular membranes. We hypothesized that like AS, molecules naturally produced by bacteria that affect membrane structure would modulate rates of vesicle exchange. For example Pseudomonas quinolone signal (PQS) has been shown to induce membrane curvature in both *Pseudomonas aeruginosa* and red blood cells and influenced vesicle production (11, 14). Many other membrane-binding molecules released by bacterial cells have been characterized, including several molecules known to have antibiotic properties. Like PQS, the membrane binding antibiotic compounds colistin and polymyxin B (PMB) increased the rate of vesicle production by bacteria (35).

Here we measured the influence of membrane-structuring molecules such as colistin, nisin, PMB and PQS on horizontal gene transfer (HGT) via EVs, as each of these molecules is known to bind to bacterial membranes, and modulation of membrane shape was been observed (45-48). Among these, colistin and PMB are known inhibitors of *E. coli* growth. In our tests, concentrations below the reported MIC were used (49, 50), see Table S2. In Table S2 we define the baseline or 1X concentration used for each compound tested. As shown in Fig S3A, colistin and PMB at this 1X concentration had a temporary effect on cell growth, although normal growth resumed after a few hours.

Colistin and PMB increased the number of cells in the population with compromised membranes, as measured using propidium iodide, but that effect was also transient as shown in Fig S3B and S3C. PQS does not have MIC reported for *E. coli* cultures and nisin does not have a well-defined MIC for *E. coli*. Their respective 1X concentrations were arbitrarily fixed at 10 µg/ml and 20 µg/ml (Table S2). We observed no decrease in the growth rate or loss of membrane integrity when *E. coli* cultures were treated either with nisin or with PQS, at 1X concentrations Fig S3. EV production and uptake were measured in the presence of each compound using the assays described in Fig. 1A. *E. coli* cells were treated with 0.5 μg/mL Bovine serum albumin (BSA) or 1 μM AHL *N*-butyryl-L-homoserine lactone (C4-AHL) were run as negative controls. Both BSA and C4-AHL are not known to bind to restructure bacterial membrane, and C4-AHL has been shown not to influence vesicle production in bacteria (51).

Vesicle production was measured by quantifying the abundance of outer-membrane proteins in purified vesicle on SDS-PAGE gel. These measurements were also compared to nanoparticle tracking analysis which directly counts EVs in solution, Fig S4. As shown in Fig 2A, Fig S5, all the three antibiotic compounds and PQS positive control increased vesicle production of the *E. coli* donor strain, similar to previous reports (11, 14, 52). Vesicle production in the presence of these compounds was concentration dependent, Fig 2B. Even upon treatment with 0.25X relative concentration, a nearly 2-fold increase in vesicle production was observed, demonstrating that even low concentrations, far below the MIC of colistin and PMB, these compounds have the potential to influence vesicle production. Vesicle size and morphology were not strongly affected by these compounds, Fig S6.

**Figure 2:**
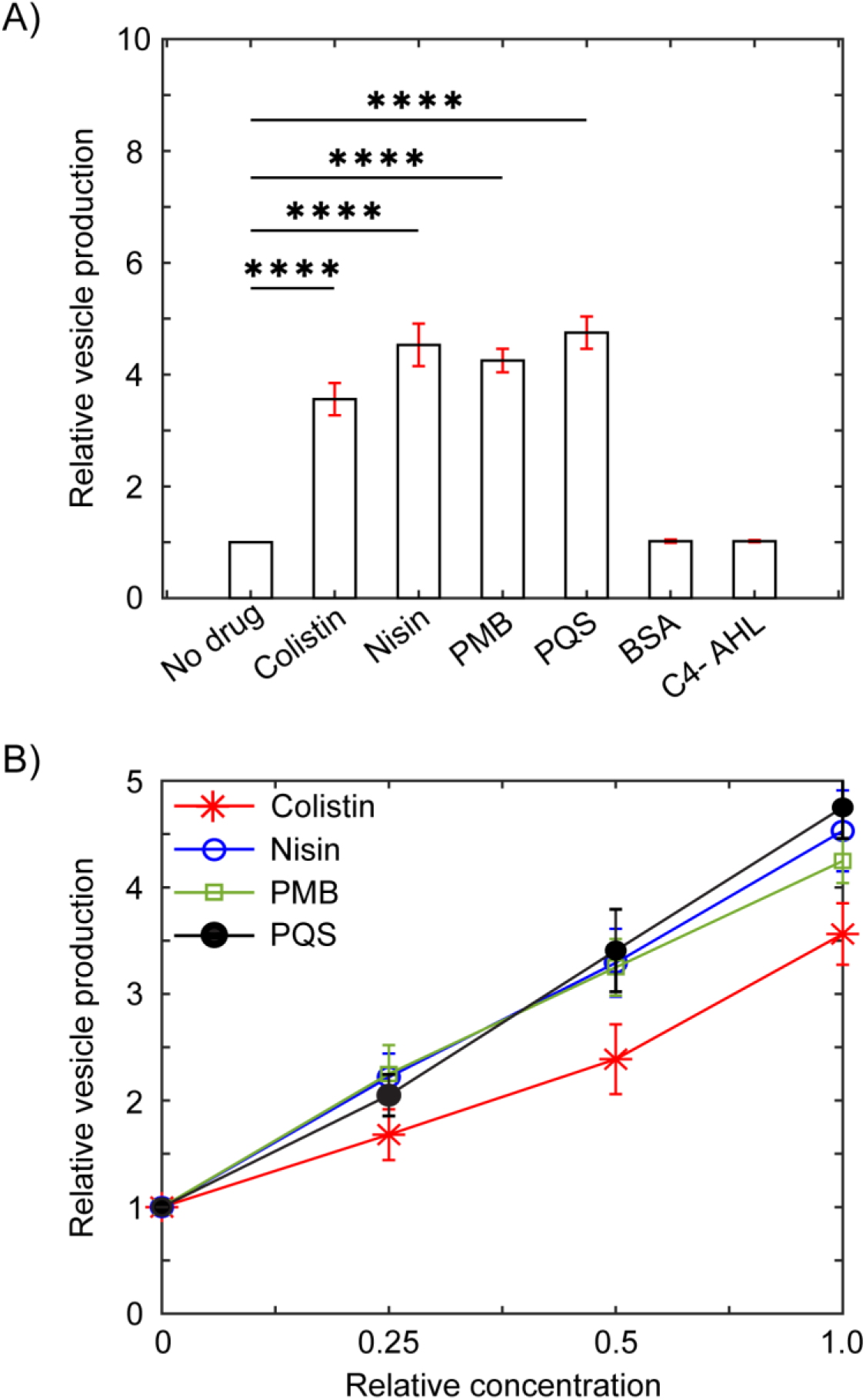
Membrane-binding compounds produced by bacteria increased vesicle production. EV production was measured by analyzing the concentration of characteristic outer membrane proteins (OmpC/F) in harvested EVs. A) Addition of exogenous molecules increased EV production in a culture of *E. coli*. (B) Vesicle production increased linearly with increase in drug concentration. 1X relative concentration for colistin and PMB is 1 µg/ml and for nisin and PQS is 10 and 20 µg/ml respectively. Error bars show standard deviation. Difference between the experimental conditions was validated with unpaired t-test (**** P ≤ 0.0001).

### Bacterial membrane binding compounds increase vesicle uptake in recipient cells

The induction of membrane curvature is also essential for vesicle fusion and therefore vesicle uptake with recipient cells. As shown in Fig. 1, alpha-synuclein, a molecule known to restructure membranes, influenced vesicle production and uptake.

Next, we tested if the four compounds shown to induce vesicle production also increased vesicle uptake. As in Fig. 1, vesicles were harvested from a donor *E. coli* strain containing the plasmid pLC-RK2, which confers kanamycin resistance to the host cells (10). Donor cells were grown in the absence of the membrane-binding compound, although as shown in Fig S7, EV transfer time was not dependent on whether EVs were produced in the presence or absence of membrane-binding compounds. Recipient cells grown to exponential phase were treated for 1 hour with one of the membrane-binding compounds prior to the addition of harvested vesicles. Cells were plated every hour on LB plates with kanamycin to track plasmid transfer. Vesicles harvested from a donor containing pLC-RK2 transferred around 10 hours in the absence of added compound.

Uptake in the presence of the 4 membrane-binding molecules tested decreased in transfer time to 5-6 hours, see Fig. 3A. Negative controls showed that 0.5 μg/mL BSA and 1 μM C4-AHL did not alter the transfer time of the plasmid. The reduction in the uptake time was dependent on the concentration of the added compound, as shown for the case of nisin in Fig. 3B.

**Figure 3.**
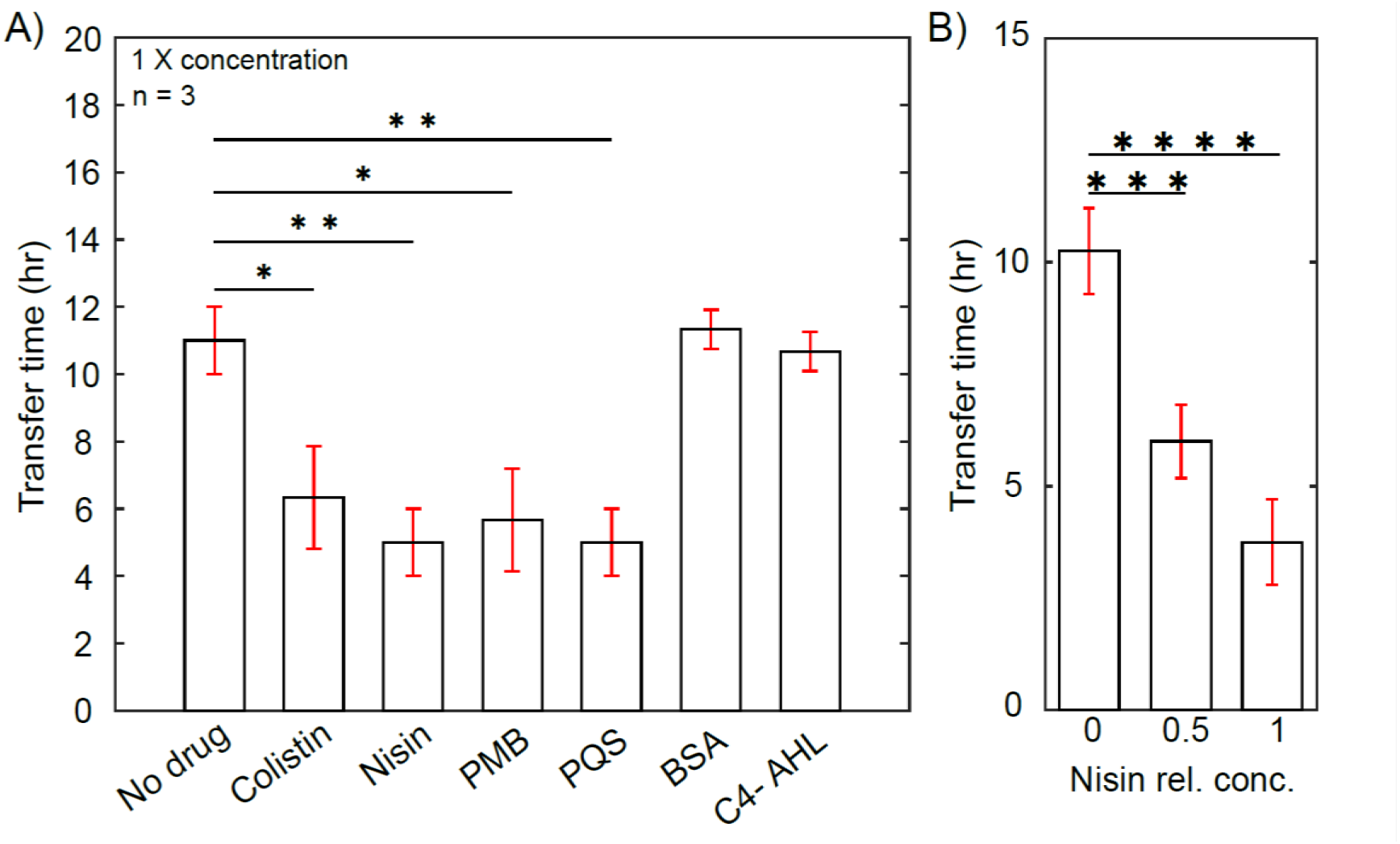
Membrane-binding compounds produced by bacteria increase vesicle uptake. Vesicle uptake was quantified as the time needed for recipient cells to gain antibiotic resistance as the result of plasmid transfer via EV uptake. A) Colistin, nisin, polymyxin B(PMB), and Pseudomonas quinolone signal (PQS) signal all increased EV uptake in a culture of *E. coli*. Bovine serum albumin (BSA) and N-butyryl-L-Homoserine lactone (C4-AHL) were negative controls. (B) nisin increased EV uptake in a dose dependent manner. Error bars show standard deviation. Unpaired t test was used to confirm the difference between treated and untreated cultures (** P ≤ 0.01, *** P ≤ 0.001, **** P ≤ 0.0001).

### Membrane binding compounds increased the rate of horizontal gene transfer within a bacterial coculture

Given that the EMs tested increase both vesicle production and uptake rates, we next tested if the addition of these compounds would regulate plasmid exchange within a bacterial coculture. As shown in Fig. 4A, exponential cultures of *E. coli* strains carrying different plasmids were mixed together. One strain was *E. coli* MG1655 carrying the pLC-RK2 plasmid with kanamycin resistance, and the other strain was *E. coli* DH5α carrying pSC101+ plasmid with ampicillin resistance. Control experiments confirmed that the plasmids were compatible and could be stably maintained in the same cell (data not shown). We hypothesized that plasmid exchange within EVs would result in a strain with resistance to both antibiotics. Strain DH5α was chosen because its genome contains a deletion of *lacZ*, enabling discrimination of the direction of gene flow via selection on MacConkey agar plates, see SI, Fig S8.

**Figure 4.**
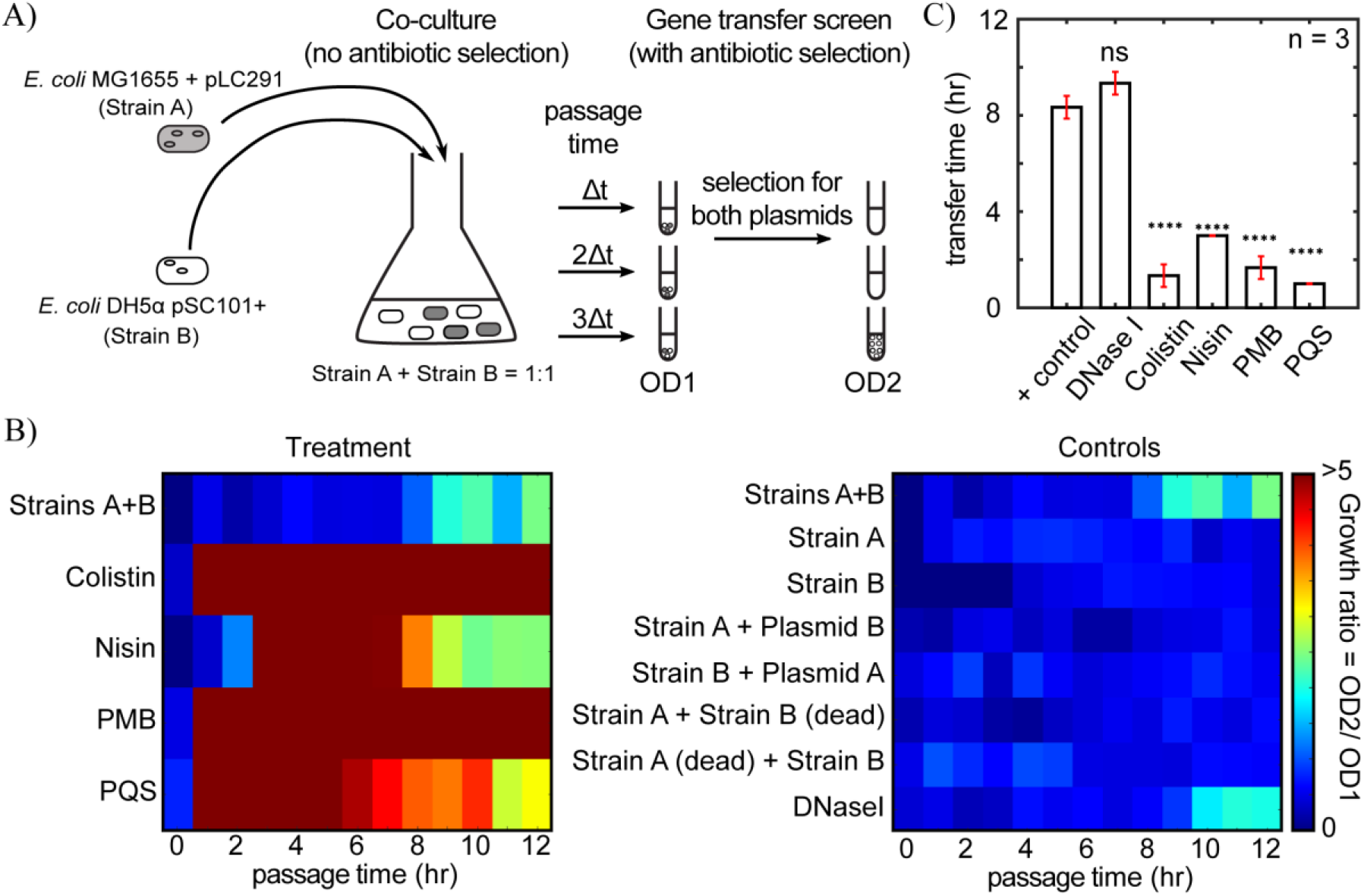
Membrane-binding molecules increased the rate of horizontal gene transfer within a bacterial coculture. A) Two strains of *E. coli* harboring plasmids with different antibiotic resistance genes were cocultured. Over time, aliquots of the co-culture were used to inoculate media containing both antibiotics to screen for cell containing both plasmids. OD1 is the optical density of cells at the beginning of the gene transfer screen and OD2 is the optical density of cells growing in double antibiotic selection after 12 hr. Growth within the gene transfer screen indicates plasmid exchange within the co-culture prior to the time of passage. B) The change in optical density within the gene transfer screen was used to compare the rate of plasmid exchange under a variety of conditions, with Strains A + B indicating the positive control. Treatments include the addition of colistin, nisin, PMB, and PQS at the 1X concentration to the co-culture. Controls include monocultures, monocultures with free plasmid, mixtures of live and dead strains, and coculture in the presence of DNaseI. C) Transfer times for the positive control, strains A + strain B, as compared to treatments with membrane binding compounds and the DNaseI negative control. Error bars indicate standard deviation Difference between the experimental conditions was validated with unpaired t-test (**** P ≤ 0.0001).

After inoculating the co-culture, 1 ml aliquots were removed every hour and used to inoculate fresh media with double antibiotic selection. This culture, called the gene transfer screen in Fig 4a, contained kanamycin at 50 µg/ml and ampicillin at 100 µg/ml, therefore only cells containing both resistance markers would proliferate. The fold change in the optical density at 600 nm after 12 hours in the gene transfer screen was used to determine whether plasmid exchange had occurred within the initial co-culture prior to the time of cell passage. As shown in Fig. 4B, in the absence of externally added membrane binding molecules (condition Strains A+B), cells with double antibiotic resistance were detected after 9 hours of coculture. For aliquots of co-culture sampled prior to 9 hours, the optical density of the culture in the presence of both antibiotics decreased over time, whereas co-culture aliquots taken at 9 hours or later resulted in an increase in optical density over time. Growth in the gene transfer screen indicated that the co-culture contained cells with both plasmids at the time of passage. Cells growing within the gene transfer screen were streaked out to form single colonies on McConkey’s agar plates with kanamycin and ampicillin, as shown in Fig. S9. PCR reactions confirmed that cell within the colonies contained both resistance genes, as shown in Fig. S10.

As shown in Fig 4B and C, in the presence of membrane binding compounds, the time needed to observe a strain with double antibiotic resistance was decreased from 8 hours to less than 4 hours. Control experiments confirmed that 1) monocultures of cells with only one plasmid did not gain double antibiotic resistance, 2) free plasmid added to a monoculture did not result in gene transfer, and 3) dead cells were incapable of transferring a plasmid. DNaseI activity within the co-culture also did not significantly change the time needed for gene transfer, suggesting that the transferred plasmid was protected from DNA degradation. Plasmid transfer within the coculture was also faster when treatments were added at 0.25 X concentration, as shown in Fig. S11.

## Discussion

Vesicle exchange is critical to many bacterial processes such as host invasion, signal exchange, and gene transfer. Vesicle exchange appears to be ubiquitous and is not known to require specialized molecular machinery for vesicle production or uptake. It seems likely there would be selection pressure to evolve strategies to regulate vesicle exchange given its potential to facilitate horizontal gene transfer. The ability of cells to regulate rates of horizontal gene transfer has been observed previously for other mechanisms of gene transfer (53-57). For example, transformation scales with the availability of free DNA. Studies have shown that killing neighboring cells increases the concentration of extracellular DNA and also the rate of transformation (58). Previous studies have revealed several biological parameters that influence vesicle production, including modulating membrane composition (13, 59), activation of stress response pathways (35, 60, 61), destruction of the cell wall, and the production of membrane structuring molecules (13, 35, 59, 60, 62, 63). It is not surprising that membrane-binding molecules would influence the production of vesicles, as eukaryotic cell utilize molecules which wedge, crowd, and bend the membrane to overcome the energetic costs of vesicle production. Here we showed that membrane-binding molecules produced by bacteria also increased the rate of vesicle uptake. Some short peptides facilitate membrane fusion, including fusion peptides and also some antimicrobial peptides (64). Membrane fusion is promoted through a combination of induction of membrane curvature, charge screening, anchoring two membranes in juxtaposition, and even modulation of membrane rupture tension (64). It remains unclear how the bacterial peptides tested here facilitate membrane fusion.

Here we tested the ability of four bacterial compounds, nisin, colistin, PQS, and PMB, to regulate vesicle exchange. Extensive work on PQS and vesicles has shown the ability of PQS to induce curvature in membranes through a wedging mechanism, which increased vesicle production (11, 65). The other compounds are classified as antibiotics, which is not surprising given that the mode of action for a large number of antibiotics is to compromise the bacterial membrane. At high concentrations these compounds coat the cell membrane, eventually forming pores that lead to cell death (66). At low, sub-inhibitory concentrations, these compounds have secondary functions, including the regulation of vesicle exchange. Pore formation does not occur at low concentrations of these molecules (67, 68), instead binding of these compounds leads to membrane bending and bleb formation, processes known to facilitate vesicle formation (52). Colistin and PMB were previously shown to induce EV formation, although the previous study focused on the ability of EVs to protect bacteria from membrane-targeting antibiotic compounds and phage infection (35). This is not the first time secondary functions have been identified for antibiotic compounds at sub-lethal concentrations (68). Low concentrations of fluoroquinolones increased conjugation (69), and sub-inhibitory concentrations of many antibiotics also act as signaling molecules (67-70). Here we show that regulation of vesicle exchange, and the associated horizontal gene transfer, is yet another secondary function of some antibiotic compounds.

There are many such membrane binding antibiotic compounds, including colistin, nisin, and PMB, and the ability of these compounds to influence vesicle production and uptake does not seem to require a specialized interaction with membrane components. An unknown component found in the supernatants of *E. coli* and *K. pneumoniae* have been shown to increase EV production in *P. aeruginosa* (5). Sub-inhibitory concentrations of gentamicin also destabilize the membrane and induce vesicle formation in *P. aeruginosa* (71). As shown here, even the eukaryotic compound AS that is involved in membrane restructuring in neurons has the ability to influence vesicle exchange in bacteria.

Therefore, many if not all membrane binding compounds, including other amphipathic alpha-helices, may have a regulatory influence on vesicle exchange. Membrane binding peptides such as antimicrobial peptides are produced by many species of bacteria, suggesting many bacteria have the potential to regulate vesicle exchange.

Recently several mechanisms of vesicle production have been reported (16, 72). Blebbing is one such mechanism, and the compounds tested here may act through this pathway given their ability to induce curvature in bacterial membranes (14, 35). Others have recently speculated that vesicles loaded by DNA are likely the result of cell explosion (16). It is possible that low concentration of membrane binding antibiotics contributes to vesicle formation through cell lysis, although it seems unlikely cell lysis accounts for the increased rate of vesicle uptake. Although many recent studies have focused on vesicle production, little work has been done on vesicle uptake by bacteria. The uptake process is essential for the transfer of biomolecules in vesicle, such as genetic material, membrane proteins, regulatory RNAs, and molecules that mediate host-bacterial interactions such as LPS (59, 73). Here we showed that compounds known to restructure the membrane facilitated vesicle uptake, but other mechanisms might also regulate the vesicle uptake rate. In eukaryotic membranes protein-protein attachments, such as SNARE proteins, are a first step in endocytosis (29). Some proteins on vesicle surfaces even insert into the membrane of recipient cells (74).

Recent work suggests uptake of bacterial vesicles into eukaryotic host cells appears to be rapid (56), although early studies on vesicle uptake via bacteria suggest uptake is a rare event (75). A better understanding of vesicle uptake and the strategies that bacteria have evolved to increase the rate of specificity of vesicle uptake, in addition to the release of membrane structuring molecules, would lead to a better understanding of vesicle exchange and its regulation within bacterial populations.

## Materials and Methods

### Bacterial Strains and Growth Conditions

*E. coli* lab strain MG1655 was used for all extracellular vesicle and transfer experiments. DH5α was also used in coculture experiments. Bacteria were grown in Luria-Bertani (LB) broth (Difco, Sparks, MD) at 37°C with shaking at 200 rpm. Plasmids were introduced to donor strains via electroporation. Plasmids were maintained in liquid culture with the appropriate antibiotics (VWR, Radnor, PA). List of plasmids are in Table S1.

### Isolation and purification of EVs

EVs were isolated from liquid cultures of *E. coli* MG1655 as previously described (10) with some modifications. 400 μL of overnight culture was used to inoculate 400 mL of LB broth containing selective antibiotic and added exogenous molecule concentration when stated. Liquid cultures were grown at 37°C with shaking at 200 rpm for 16-20 h. Cells were pelleted by centrifugation at 1,200 x g at 4°C for 30 min. The supernatants were decanted, and vacuum filtrated through ExpressPlus 0.22 μm pore-size polyethersulfone (PES) bottle top filter (Millipore, Billerica, MA) to remove remaining cells and cellular debris. Vesicles were collected by ultra-centrifugation at 80,000 x g (Ti 45 rotor; Beckman Instruments, Inc., Fullerton, CA) at 4°C for 1.5-2h followed by 180,000 x g (Ti 70i rotor; Beckman Instruments, Inc., Fullerton, CA) at 4°C for 1.5-2h and resuspended in 1mL of phosphate buffered saline (PBS) and stored at 4°C. Vesicle preparations were treated with 100 ng mL^-1^ of DNase I at 37°C for 20 min followed by deactivation of the DNaseI at 80°C for 10 min. Vesicle preparations were also plated on LB agar to check for the presence of bacterial cells.

### EV quantification

Extracellular vesicle concentrations were quantified using SDS-Polyacrylamide gel electrophoresis. Vesicle preparations were treated with 6xSDS loading buffer and boiled for 10 min at 100°C and run on a 10% SDS-PAGE gel (Bio-Rad Laboratories, Hercules, CA), stained for 15 min with Coomassie Brilliant Blue Stain, and destained in H_2_O, methanol, and acetic acid (50/40/10 v/v/v) overnight. Protein concentrations of OmpC/F were determined using ImageJ from a standard curve generated by a BSA protein concentration gradient, as shown in Fig. S1. Protein concentrations of OmpC/F were used to quantify vesicle concentration and production relative to untreated cells.

### Exogenous molecules used

Colistin sulfate salt, nisin, polymyxin B sulfate and 2-Heptyl-3-hydroxy-4(1H)-quinolone (PQS) (Sigma-Aldrich Corp., St. Louis, MO) were dissolved in water. Purified alpha-synuclein was provided by Ralf Langen’s lab at USC (32).

### Measurement of bacterial growth

Overnight grown culture of *E. coli* MG1655 was used to start parallel cultures treated either with 1 µg/ml colistin or with 10 µg/ml nisin or with 1 µg/ml PMB or with 20 µg/ml PQS, at 1% inoculum. The growth of all cultures was monitored in 96-well at 600 nm using plate reader (TECAN, infinite M200PRO) for 12 hrs at 37°C with intermittent shaking for 30 secs.

### Propidium iodide assay

25 ml Secondary cultures of *E. coli* MG1655 were grown at 37°C, 200 rpm till OD reach ∼ 0.2, after which individual cultures were subjected to the treatment with 1 µg/ml colistin or with 10 µg/ml nisin or with 1 µg/ml PMB or with 20 µg/ml PQS. Treated cells were harvested at 0^th^, 2^nd^, 5^th^ and 10^th^ hours, washed thrice with 1X PBS and stained with Propidium Iodide Ready Flow™ Reagent (Invitrogen by Thermo Fisher Scientific) at 25°C. Culture were again washed with 1X PBS and fixed with 4% PFA. 5 µl of aliquot from these cultures was then spread on the slide and imaged with 40X/ 0.6 NA objective on ECHO revolve microscope in Phase contrast and RFP channel.

### EV-mediated gene transfer

Gene transfer experiments were modified from previously published work (10). The *E. coli* recipient strain was diluted 1:1000 from overnight culture in 4 mL LB broth (Difco, Sparks, MD) at 37°C with shaking at 200 rpm to early log phase, OD_600_ 0.2, ∼2 h, and exogenous molecules were added and incubated for 30 mins. Then at time 0 h, purified vesicles were added. The number of vesicles added to recipient cultures was standardized for transfer experiments. In all transfer assays, vesicles equivalent to 1 µg of the outer membrane proteins OmpC/F were used. Every hour, 200 μL of culture was removed and plated on LB agar plates containing either 50 μg mL^-1^ kanamycin or 50 μg mL^-1^ carbenicillin or both dependent on plasmid resistance. After 16 h of incubation at 37°C, plates were counted and scored for CFUs. The bacterial colonies that acquired antibiotic resistance were re-selected on antibiotic selection plates and the presence of the transferred plasmid was verified for several colonies using PCR. Gain of resistance not associated with plasmid transfer was not observed.

### EV coculture gene transfer

Coculture experiments were performed using DH5α (Δ*lacZ*) cells transformed with pSC101+ (*bla*) and MG1655 (with *lacZ)* transformed with pLC-RK2 (*npr*). Each strain was grown separately starting in overnight cultures and mixed together in 1 : 1 proportion next day. The resultant inoculum was added to fresh 100 ml LB at 1% inoculum and grown at 37°C with shaking at 200 rpm for another 12 hours. 1 ml samples withdrawn periodically after every hour from the coculture were washed thrice with 1X PBS and inoculated in 10 ml LB containing ampicillin (100 µg/ml) and kanamycin (50 µg/ml). Optical densities of these cultures were recorded for all 13 time points (*SPECTRONIC 200*, Thermo Fisher Scientific) and denoted as OD1. This was followed by incubation of cultures at 37°C and 200 rpm approximately for 12 hrs and measured for changes in respective optical densities (OD2). Ratio of OD2 to OD1 was used to determine if growth occurred in the presence of selection for both resistance markers. Growth indicated the presence of cells harboring both plasmids, as confirmed by PCR (Fig S10). Cultures at time points with ratio above 1 were streaked on MacConkey’s agar (Sigma-Aldrich) containing ampicillin and kanamycin to differentiate between the two hosts.

### Transmission electron microscopy

Transmission electron microscopy specimens were prepared on carbon-coated formvar grids (Electron Microscopy Sciences, Hatfield, PA). Samples were absorbed on the grids for 5 m and then negatively stained with 1% (w/v) aqueous uranyl acetate. Images were taken on a JEOL 1400 transmission electron microscope (JEOL USA Inc., Peabody, MA) at an accelerating voltage of 100kV.

### Nanoparticle tracking analysis (NTA)

Malvern Panalytical Nanosight NS300 was used (Malvern, UK) equipped with a 532 nm green laser. All samples were diluted in PBS to a final volume of 1 mL. For each measurement, five 1-min videos were captured with detection threshold of 5, embedded laser, 45 mW. After capture, videos were analysed by the in-build Nanosight Software NTA 3.1 Build 3.1.46.

### Colony PCR

PCR was performed using colonies from transfer assays using One*Taq* (New England BioLabs Inc., Ipswich, MA). Briefly, the reaction mixtures consisted of 0.5 μl of bacterial colony resuspended in H2O, 0.2 μM primers, and 1 U of One*Taq* polymerase (New England BioLabs Inc., Ipswich, MA) in a final volume of 25 μl. The program consisted of 25 cycles of denaturing at 94°C for 30 s, annealing at 60°C for 60 s, and extension at 68°C for 30 s. Primers used for pLC-RK2 were: forward-5’ CATTCGTGATTGCGCCTGAG 3’; reverse- 5’ TCAACGGGAAACGTCTTGCT 3’; pSC101: forward- 5’ AGTGATAACACTGCGGCCAA 3’; reverse- 5’ TGAGGCACCTATCTCAGCGA 3’.

## Acknowledgements

We would like to thank Ralf Langen’s lab for providing the purified sample of alpha-synuclein and for help with Transmission electron microscopy, Paolo Neviani and the ExtraCellular Vesicle Core at Children’s Hospital Los Angeles for help with nanoparticle analysis, and Carla Vidal Silva for helpful discussion and preliminary work on this project. JB and FT acknowledge funding from NSF award MCB-1818341, NSF award PHY-1753268, Army Research Office MURI Award W911NF1910269.

